# *Leishmania amazonensis* infection induces IL1β-dependent hyperalgesia, while dampening mechanical allodynia in C57BL/6 mice

**DOI:** 10.1101/2025.02.18.638800

**Authors:** Gustavo G. Perez, Emanuelle V. de Lima, Leticia S. Antonio, Eduardo Matheus M. Vidal, Raissa R. Christoff, Milla B. Paiva, Antônio José S. Gonçalves, Marcia Pereira O. Duarte, Maria Bellio, Robson Coutinho Silva, Robson da Costa, Rodrigo T. Figueiredo, Julia R. Clarke, Herbert L. Guedes

## Abstract

Leishmaniases are neglected diseases causing significant deaths and disabilities. In Brazil, the most prevalent form is cutaneous leishmaniasis, characterized by painless lesions despite intense inflammation and ulceration. While BALB/c mice models exhibit hypersensitization to inflammatory stimuli, C57BL/6 mice better mimic human-like lesion progression and nociceptive responses. This study aimed to investigate the mechanisms underlying nociceptive changes in cutaneous leishmaniasis using the C57BL/6 model. Following infection with *L. amazonensis*, behavioral and nociceptive tests revealed unaltered mechanical nociception and motor capacity, though thermal hypersensitivity emerged during the chronic phase. Elevated IL-1β production in the lesions and upregulation of TRPV1 in dorsal root ganglia (DRG) neurons were detected via ELISA and qPCR. Mice deficient in IL-1β-related proteins or receptors exhibited higher thermal nociception thresholds, highlighting IL-1β’s role in heat hypersensitization during late infection stages. Microscopy of chronic lesions revealed tissue deformities, indicating desensitization to mechanical and inflammatory stimuli due to nerve terminal alterations and fibroplasia from regenerative processes. Conversely, thermal hypersensitivity in chronic phases was driven by IL-1β effects on thermal nociceptive neurons in the DRG. These findings suggest that IL-1β and TRPV1 contribute to thermal hypersensitivity, while structural changes in lesions underlie mechanical desensitization. This model provides insights into the complex nociceptive mechanisms of cutaneous leishmaniasis.

## 1. Introduction

Leishmaniases are a group of neglected tropical diseases caused by parasites of the genus *Leishmania* with higher impact on developing countries. In Brazil, leishmaniasis is endemic, and affects thousands of people every year, causing humanitarian and economical losses^1,2^. Localized cutaneous leishmaniasis (LCL) is the most prevalent clinical manifestation of this disease and is most frequently associated with infections by *Leishmania amazonensis* (*L. amazonensis*), *L. braziliensis* and *L. guyanensis* ^3^.

Numerous reports of different forms of leishmaniasis causing neurological symptoms have been made, either in patients or other animal models^4–6^. The presence of painless ulcers in LCL, despite intense inflammatory response, may be related to commitment of the nervous system, which has been found previously in patients and other animal models^7,8^, or to pain-reducing metabolomic reprogramming^9^. Among those, peripheral nerve damage^10^ and even the presence of parasites in the Central Nervous System ^11^ have been reported. However, there is no consensus about whether the infection is painful or painless^12^, especially in mice, as there is data supporting both hypotheses^9,13,14^. For example, mice with different backgrounds seem to differ in response to noxious stimuli. BALB/c mice infected with either *L. major* or *L. amazonensis* showed early phase hyperalgesia, which may remain until later stages^15,16^. However, there is also data providing evidence for hypoalgesia, especially at later stages of the disease^13^. Conversely, C57BL/6 mice showed thermal hyperalgesia in the same study, however, studies with these mice remain mostly unexplored in the context of pain in LCL. BALB/c mice are canonically more susceptible to the infection because of the predominant T helper 2 (Th2)-mediated immunological response leading to anti-inflammatory response along with macrophage repair environment^17^. In contrast, the C57BL/6 lineage adopts a resistant phenotype, with more similarities to what happens in patients, due to its ability to induce inflammation and a proinflammatory environment for macrophages^18^.

One of the main events underlying the hyperalgesia-inducing response in *Leishmania* infection is the production of IL-1β due to the inflammasome^19–21^. The inflammasome is a protein complex that may be formed by different adaptor proteins, such as the NLRP3, and when fully formed, it leads to Caspase-1 activation, cleaving the inactive pro-IL-1β into its active form^22^. Caspase-11, in its turn, is responsible for the activation of the non-canonical pathway of inflammasome activation pathway, also culminating in Caspase 1 activation^22^.

In this study, we investigated whether the infection by *L. amazonensis* is able to induce pain or desensitization in C57BL/6 mice, and if so, which are the mechanisms and molecular pathways underlying these alterations. We observed two separate phenomena occurring: peripheral desensitization to prohyperalgesic inflammatory stimuli due to nerve terminals damage, leading to the absence of inflammation-induced mechanical allodynia; and central sensitization of non-responsive nociceptors to the mechanical stimuli due to interleukin-1β (IL-1β) release and interaction with its receptor IL-1R1 in TRPV1^+^ bearing neurons.

## 2. Materials and Methods

### 2.1. Experimental design

In total, three independent experiments were performed, and data obtained from one of these replicates are shown in each graph/figure. Animals were monitored throughout the infection and different phases were evaluated: peak phase (60 days post-infection (dpi)), when the infection-induced lesion reaches its maximum size; and chronic phase (from 90 dpi until 180dpi), when lesion size is either constant or reduced.

Behavioral assessments were performed before and after infection. Tissues were collected for transcription analysis (qPCR), histological analyzes (H&E) and cytokine level analysis by Enzyme-Linked Immunosorbent Assay (ELISA).

### 2.2. Animals

All procedures were previously approved by the Institutional Committee for Animal Care and Use of the Federal University of Rio de Janeiro (protocol n° 092/19 and 024/20) and followed the ARRIVE guidelines.

Female C57BL/6 mice were obtained from the Central Animal Facility of the Health Sciences Center (CCS), at UFRJ, and were 6 - 12 weeks-old at the moment of infection.

Animals were maintained under controlled conditions of temperature (22 ± 2°C), humidity (60-80%) and a 12-hour light/dark cycle. Each housing cage was used to store up to 5 animals, with free access to regular chow and water. Wild Type (WT) mice, or transgenic animals deficient for the caspase 1 and 11 genes (Casp1/11^-/-^), the NLRP3 gene (NLRP3-/-), the IL-1R gene (IL-1R^-/-^), the IL-18 gene (IL-18^-/-^) or the IL-18R gene (IL-18R^-/-^), were obtained from the Transgenic Animal Laboratory from UFRJ. Controls (Ctrl) were either mice challenged with phosphate buffered saline (PBS), or mice assayed for the contralateral.

### 2.3. Infection

*L. amazonensis* parasites (MHOM/BR/75/Josefa strain) were used in the infection. Amastigote forms were isolated from the lesion of infected BALB/c mice, maintained as promastigotes in M199 medium (Sigma-Aldrich) supplemented with 10% fetal bovine serum (SFB, Cultilab) and hemin (2 μg/ml), and incubated at 26°C.

For the infection, mice were anesthetized with isoflurane using a vaporizer system (Bonther BP-150; São Paulo, Brazil), and then injected with 2x10^6^ stationary phase promastigotes cells per milliliter in 20 μL PBS, into the intraplantar region of the right hind paw with a Hamilton syringe. Control groups received intraplantar administration of 20 μL of PBS. Lesion size was monitored weekly using a caliper (Mitutoyo, Brazil), where the diameter of the infected and contralateral paws was vertically measured. Size variation (Δ thickness) was determined by comparing each measurement of each paw with baseline value (before *L. a.* or PBS challenge). To guarantee infectivity, the parasites used were in the third culture passage or lower.

After skinning the hind paws and homogenizing in 1 ml of M199 medium using a tissue grinder, homogenates were used to determine the parasite load by being submitted to limited dilution assay (LDA).

### 2.4. Nociception and behavioral tests

All tests were preceded by a 1 h-long acclimatization stage in the same room where experiments were to be performed, under a direct source of light.

#### 2.4.1. Mechanical hypersensitivity (von Frey test)

To study whether animals developed mechanical allodynia, they were individually placed in clear Plexiglas boxes (9x7x11cm) on elevated wire-mesh platform (75x33 cm) (Ugo Basile, Comerio, VA, Italy), allowing access to the plantar surface of the paws. Animals were acclimated to the boxes for 1 h per day, for 3 consecutive days, before the test began. A series of Von Frey hair filaments (Bioseb – in vivo research instrument) with diameters ranging from 0.02 - 2.0 g were applied to both paws. Tests were initiated with the 0.6 g filament, and in the absence of a nocifensive response, incrementally stronger filaments were used consecutively until a response was elicited, as previously described^23^. The data collected using this up-down method were utilized to calculate paw withdrawal thresholds (in g). Baseline measurements were recorded prior to the injection of nociceptive stimuli. Significant reductions in paw withdrawal thresholds were considered indicative of mechanical allodynia^24^.

#### 2.4.2. Formalin test

The formalin test was performed at 180 dpi, by analyzing nociception induced by formaldehyde applied to the paw of mice. A 20 μL solution of 0.5% formaldehyde was injected into the intraplantar region of the contralateral paw and reactions indicating symptoms of hypernociception, such as licking, biting or paw shaking were timed, from 0 - 5 min (phase 1) or 15 - 30 min (phase 2) after formalin injection. Phase 1 of this test is typically associated with direct activation of nociceptors^25^, causing hypersensitization and the production of algogenic substances related to neuronal damage, such as neuropeptides and substance P. In contrast, responses seen in later stages of this test are caused by inflammatory stimuli (inflammatory pain)^26^. After this test, animals were euthanized to prevent tissue damage and additional suffering.

#### 2.4.3. Assessment of cold allodynia

Challenged mice were evaluated for cold allodynia by the acetone test. The same apparatus described in 2.4.1 was used, consisting of metal mesh platforms and Plexiglas boxes. Approximately 100 μL of acetone were instilled either in the plantar region of the challenged or contralateral hind paws, and manifestation of pain-indicative behaviors such as paw licking and shaking were monitored for 1 minute. This entire process of instillation and monitoring of behavior was repeated, and results from both tests were averaged for determination of reaction time. Latency to show signs of pain was measured using a stopwatch. A significant increase in nocifensive behaviors after cold stimulus was interpreted as indicative of cold allodynia, as described previously^27^.

#### 2.4.4. Hot plate test

The hot plate test was performed to evaluate thermal nociceptive stimulus and reaction time of the animals. Each animal was then positioned above a heated metal plate (INSIGHT, Brazil) set at 55 °C and the time it took until animals showed a reaction of discomfort for the first time (licking, shaking the paw, or jumping) was determined by an experient researcher using a stopwatch, and then they were removed from the plate. A maximum cutoff time of 15 seconds was considered for each animal, to avoid potential tissue damage or stress.

#### 2.4.5. Open field test (OFT)

This test was carried out at 60, 100 and 130 dpi to evaluate the exploratory and locomotor activity of the animals. An OF wooden box measuring 0.3 (width) x 0.3 (length) x 0.45 (height) m was used. Animals were placed in the center of the arena, under indirect light, and allowed to freely explore it for 5 minutes. The arena was thoroughly cleaned with 70% ethanol in between trials to eliminate olfactory cues. ANY-Maze software (Stoelting Company, Wood Dale, IL) was used to assess the total distance traveled and the amount of time spent by each mouse in the center of the box.

#### 2.4.6. Rotarod performance test

The test was performed in a rotarod mouse apparatus (Insight Ltda., Brazil), as previously described^28^. Briefly, mice at 130 dpi were positioned individually on the floor of the apparatus for 3 minutes, followed by a 2-minute acclimatization session on the rotating cylinder. The test was performed three times with an interval of 60 min between tests, positioning the animals again at the top of the cylinder, which had its rotating speed gradually increased (minimum speed 16 rpm, with a maximum speed of 36 rpm and acceleration rate of 3.7 rpm). Latency to fall was recorded over a 5-min period, and results were expressed as mean latency among tests.

### 2.5. Tissue collection and preparation

After 5-6 months of infection, animals were sedated with an intraperitoneal injection of ketamine and xylazine (90 mg/kg ketamine and 4.5 mg/kg xylazine, i.p) and perfused with saline through the heart for at least 2 minutes, until the liver showed discoloration. We then collected the region of the dermis and epidermis of the paws, cutting the tissue before the tendons. This tissue was processed for histological analyses. The rest of the paw was used for cytokine analysis by ELISA. Dorsal Root Ganglion (DRGs) were collected from the challenged and contralateral sides (L1 to L6 segments), and frozen immediately for qPCR analysis, while the spinal cord was collected for immunofluorescence analysis. For quantification of parasite load, the infected paw was excised and disinfected by placing it for 1 min in 70 % alcohol. Lymph nodes were also harvested and placed in Eppendorf tubes containing 1 mL of 199 medium with 10 % FBS. These organs were then homogenized. A volume of 50 μL of the homogenates was diluted four-fold in 150 μL of 199 medium per well, in 96-well plates, which were then placed in a BOD incubator for 7 to 14 days at 26 ◦C. By the end of this timeframe, plates were checked under an optical microscope and, for the determination of the parasite load, the last well in which parasites were observable was considered.

### 2.6. Hematoxylin & eosin staining and tissue visualization

This process was carried out by the Histotechnology Platform at Fiocruz-RJ, Cardoso Fontes Pavilion, in accordance with the Standard Operating Procedures: POP-PlatHisto-TEC-001/002/003/005. Briefly, after collection, tissues were fixed in 4% paraformaldehyde for 5 days, dehydrated in ethanol gradient (70%, 80%, 95%, 100%), infiltrated and embedded in paraffin. The resulting histological sections (4-5 μm thick) were stained with hematoxylin and eosin. The sections were analyzed using an AxioHome Carl Zeiss bright field microscope, with an HRc5 camera, in collaboration with the Laboratory of Experimental Medicine and Health, IOC/Fiocruz.

### 2.7. Immunohistochemistry for spinal cord

The spinal cord of C57BL/6 mice was fixed with 4% paraformaldehyde (Sigma-Aldrich, USA) overnight and then embedded in 30% sucrose to provide cryoprotection. Coronal sections (10 μm) were obtained with a cryostat (Leica, Germany) to perform immunohistochemistry. After standard antigen retrieval, slides were incubated with 0.3% Triton X-100 (Sigma-Aldrich, USA and incubated with 10% normal goat serum (Sigma-Aldrich, USA) for 30 min. Next, the following primary antibodies were incubated overnight: anti-rabbit TRPV1 (1:1000, Invitrogen, PA 129421) and anti-mouse ILR1 (1:500, Santa Cruz, sc-393998). Then samples were washed with phosphate-buffered saline (PBS) and incubated with secondary antibodies: goat anti-rabbit Alexa Fluor 488 (1:500; Abcam, ab150077), and goat anti-mouse Alexa Fluor 594 (1:500; Abcam, ab150116) for 2 hours, washed and mounted with fluoroshield with DAPI (ab104139). Images were acquired with a ZEISS confocal microscope (LSM 710).

### 2.8. Cytokine production analysis

When animals were euthanized, their paws were removed and homogenized with a tissue homogenizer (Ultra Turrax T10 Basic, IKA, Germany) in PBS. The homogenates were then centrifuged for 10 min at 2,000xg, 4 oC and the supernatants were collected and stored at -20 oC until use.

The levels of IL-1β and tumor necrosis factor α (TNF-α) were analyzed by ELISA, following the manufacturer’s instructions (BD OptEIA/R&D Systems) using the supernatant from hind paws of animals challenged with *L. amazonensis* or PBS.

### 2.9. RNA extraction from tissue and preparation of cDNA

RNA extraction was performed using TRIzol reagent, according to the manufacturer’s instructions (Invitrogen). To this end, 100 mg of tissue were macerated in 300 μL of TRIzol reagent, followed by addition of 150 μL of chloroform. After 5 minutes of incubation at room temperature, the mixture was centrifuged at 12,000 x g for 15 minutes at 4 °C, and the aqueous phase was carefully separated from the organic phase, added to 150 μL of isopropanol and left overnight at -20 °C. The pellet was then washed 3 times with 500 μL of 75% ethanol in DEPC water, dried at room temperature for 10 minutes and resuspended in DEPC water. The purity and integrity of the RNA were determined using the absorbance ratios at 280/260nm and 260/230nm in a Nanodrop 1000 spectrophotometer (ThermoFisher Scientific Inc.). Only preparations with ratios above 1.8 were used for the synthesis of complementary DNA (cDNA).

### 2.10. Gene expression analysis by qPCR of purified cDNA

One μg of total RNA was treated with DNAse I (ThermoFisher Scientific Inc) according to manufacturer’s recommendations before complementary DNA (cDNA) synthesis using the High-Capacity cDNA Reverse Transcription Kit (ThermoFisher Scientific Inc). Analyses were carried out on QuantStudioTM 5 equipment (ThermoScientific Inc, Waltham, MA) and the expressions of the genes of interest were obtained using the Syber Green Master Mix kit (ThermoFisher Scientific Inc, Waltham, MA). Cycle Threshold (Ct) values were normalized with a housekeeping gene and analysis using the ΔΔCt method to generate fold change values relative to control groups. Actin gene was used as an endogenous control and the primer sequences used are listed in Table 1.

**Table 1.**
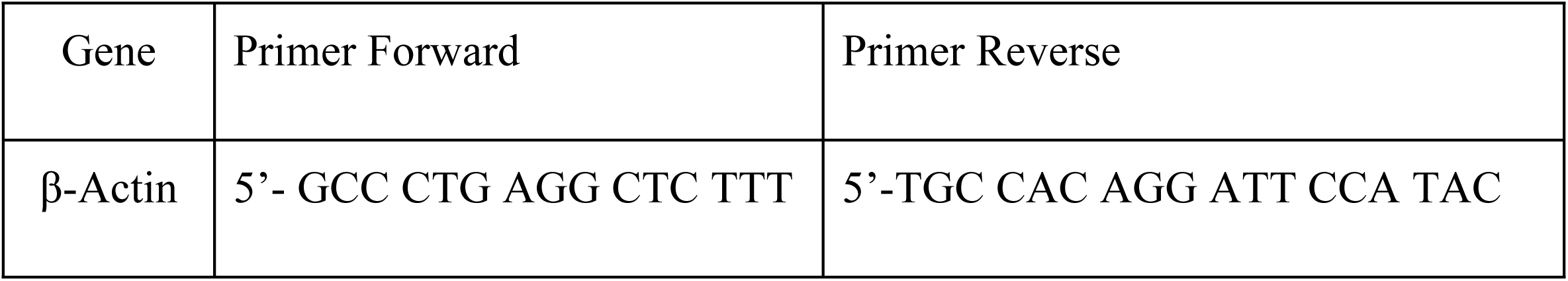
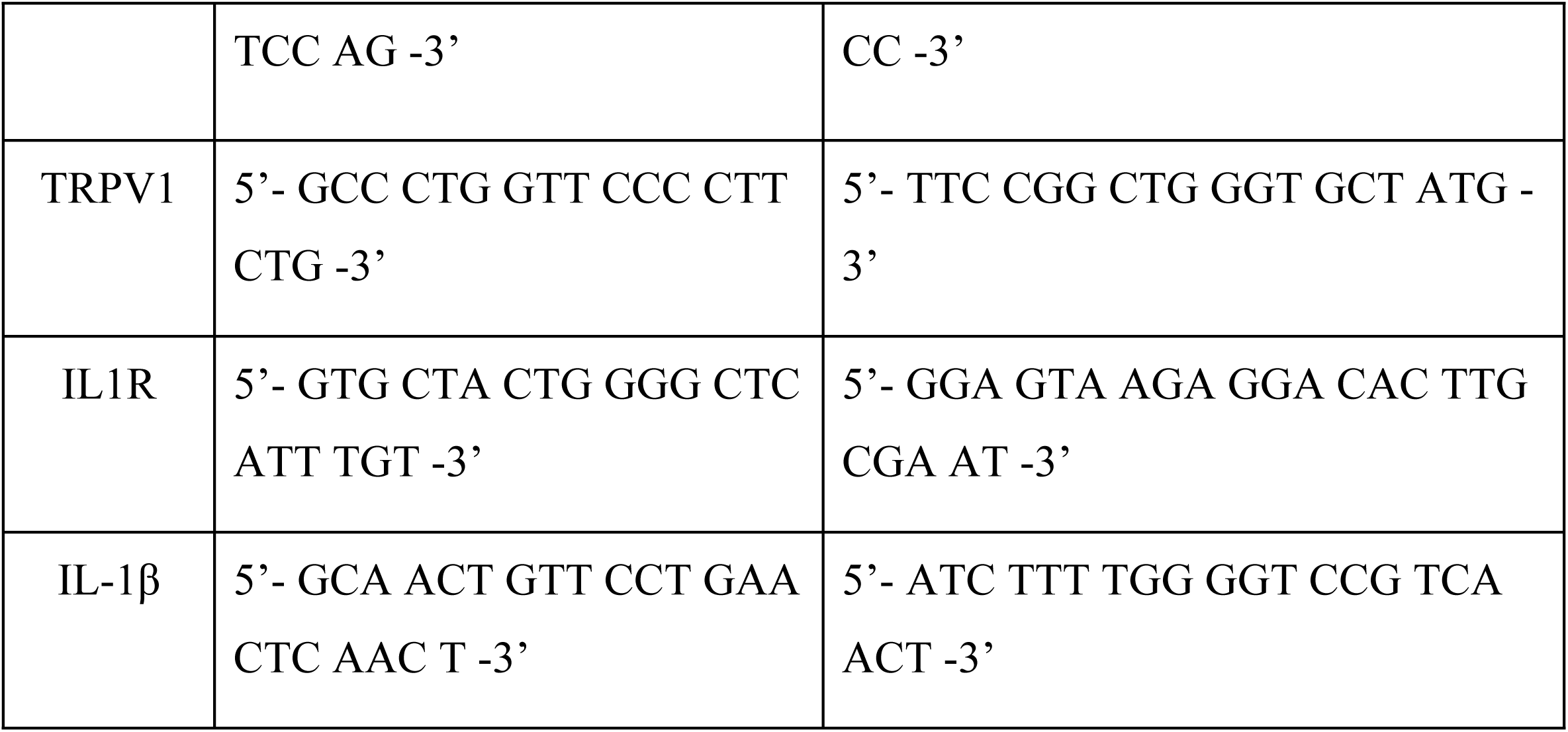
Primers used for qPCR. Forward and reverse primers designed for measuring the expression of the designated genes.

### 2.11. Statistical analysis

Data were expressed as mean ± standard error (SEM) or standard deviation (SD). Significant differences were studied using unpaired Student’s t-test when there were two groups and one time point of analysis, or analysis of variance (ANOVA) if more groups or more than one time point was analyzed. For comparison between groups, Tukey’s multiple comparison test was carried out following one-way ANOVA, or Dunnet’s multiple comparison test in two-way ANOVA. p < 0.05 were considered statistically different, and p values are indicated in the corresponding figure legend. Degrees of freedom (df) were standardized as 1 for all tests. All statistical analyzes were performed using GraphPad Prism 8®.

## 3. Results

### 3.1. The lesion caused by the infection with *L. amazonensis* does not cause mechanical allodynia in C57BL/6 mice

Inflammation is usually associated with the induction of allodynia and hyperalgesia^29^. Therefore, the inflammatory process inherent to the infection would be expected to induce hypersensitization. To keep track of the inflammatory process induced by the infection, we challenged mice in the lower right hindpaw, performed behavioral studies in the course of the infection, and weekly measurements of paw size (Suppl. Fig. 1). The values obtained by measurement in each time point were subtracted from the values before infection (Δ paw width). As expected, lesions grow until they peaked, when they stagnate and maintain their size throughout the infection (Fig. 1A). Regarding mechanical sensitivity, we performed the Von Frey test, which can measure the paw withdrawal threshold, and thus mechanical allodynia by comparing Ctrl and infected animals (Fig. 1B). Like what is observed in LCL lesions in patients, C57BL/6 did not show signs of allodynia, either in the peak phase (Fig. 1B, left, 60 dpi), nor in the chronic phase (Fig. 1B, right, 120 dpi). Together with this result, we could observe no correlation between Δ paw width and paw withdrawal threshold (Fig. 1C), further supporting that inflammation caused by the infection is not able to induce mechanical allodynia.

**Fig. 1.**
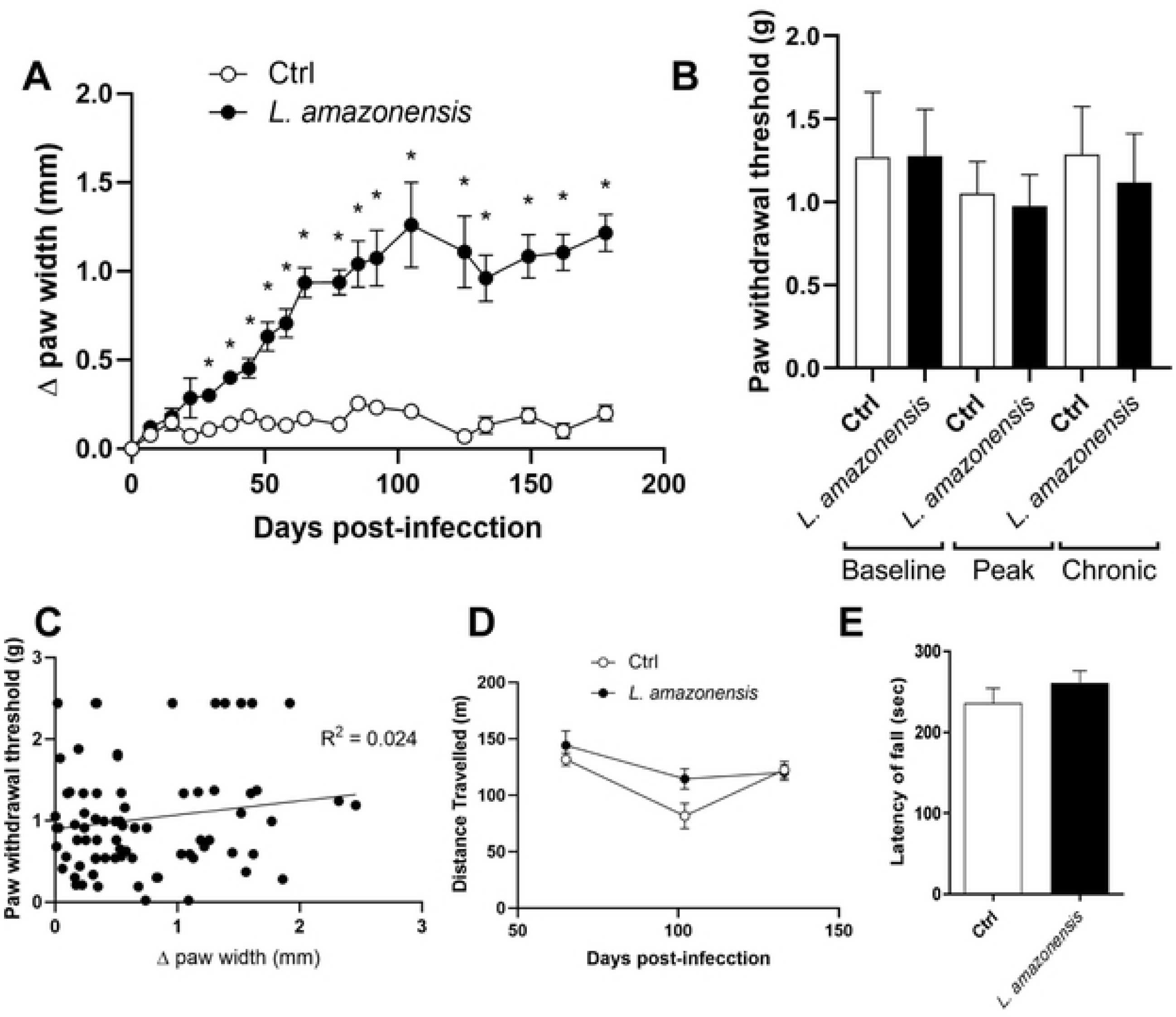
Lesion caused by the infection with *L. amazonensis* does not cause mechanical allodynia in C57BL/6 mice. (A) Adult C57BL/6 mice were challenged subcutaneously in the right hind paw with 2 x 10^6^ cells/mL promastigotes of *Leishmania amazonensis* and were assessed throughout infection. Δ paw width was measured every week throughout the infection. Data expressed by mean ± SEM, *p<0.01 by Two-way ANOVA, Sidak’s multiple comparison test, with individual variances computed for each comparison (n=10/group). (B) Mice were also evaluated according to their response threshold to the Von Frey test in the peak phase, 60 dpi, or chronic phase, 120 dpi. Data expressed by mean ± SEM, One-way ANOVA, Tukey’s multiple comparison test, with a single pooled variance (n=6/group). (C) Correlation between response threshold and Δ paw width, each data point correlates to a single response in each day throughout the whole experiment, correlation tests pointed out negative results (R²= 0.024). Data expressed by individual values, and linear regression. (D) Distance traveled in the OFT. Data expressed by Two-way ANOVA, Sidak’s multiple comparison test, with individual variances computed for each comparison (n=4-10/group). (E) Latency to fall in the rotarod test. Data expressed by mean ± SEM, t-test, p=0.342 (n=4-5/group).

Aiming to confirm whether the lesion caused by *L. amazonensis* infection was inflicting any nuisance on the mice, we conducted behavioral analysis for locomotion with the OFT (Fig. 1D). This test also allows us to measure anxious-like behavior through comparison of time spent in the center/periphery, but we could see no difference (Data not shown). Along with that, motor ability was assayed through latency of fall in the Rotarod test (Fig. 1F). There was no evidence for change in locomotion or motor activity.

### 3.2. Radical changes in the infected tissue and nerve terminals are caused by the infection with *L. amazonensis*

Aiming to understand possible changes responsible for the dampening of mechanical allodynia, we did microscopical evaluations of the skin in the challenged paws 180 dpi. Skin from mice challenged with *L. amazonensis* showed changes, such as thickening of the epidermis, with multiple layers of keratinocytes when compared to that of Ctrl (Fig. 2A,B). Besides, intense inflammatory infiltrate was seen in both the dermis (Fig. 2C) and epidermis (Fig. 2D), with parasite-bearing mononuclear cells, and even amastigotes not associated with any cell.

**Fig. 2.**
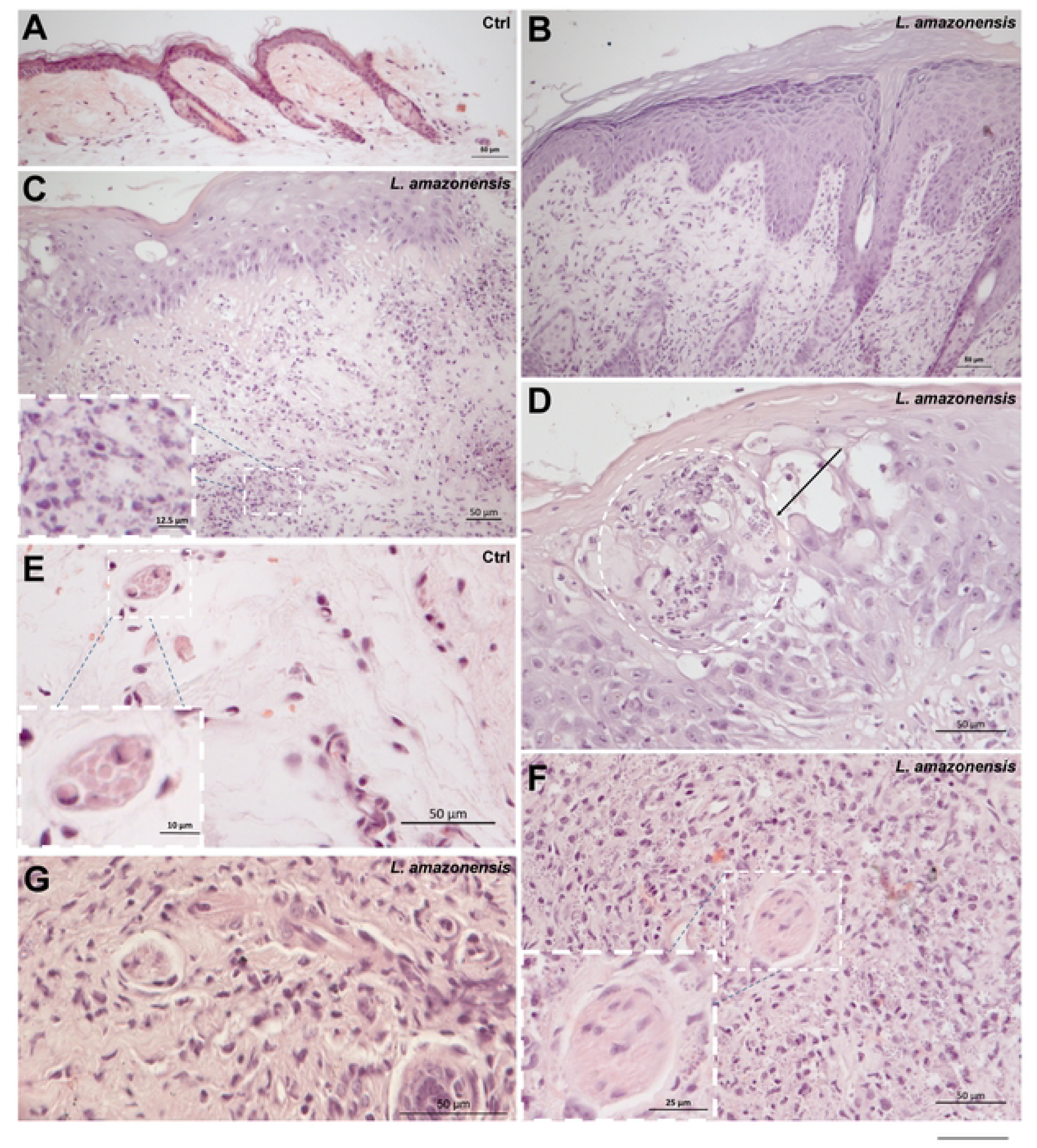
Radical changes in the infected tissue and nerve terminals are caused by the infection with *L. amazonensis*. Adult C57BL/6 mice were challenged subcutaneously in the right hind paw with 2 x 10^6^ cells/mL promastigotes of *Leishmania amazonensis* and tissue was collected 6 months after challenge. Representative images of H&E staining, with scale bars represented in the lower right. (A) Epidermis of Ctrl mice. Scale bar: 50 μm. (B) Thickening of the epidermis of challenged mice. Scale bar: 50 μm. (C) Dermis of challenged mice showing presence of *Leishmania* amastigotes. Scale bar: 50 μm. Inset: amastigotes associated or not with host cells. Scale bar: 12.5 μm. (D) Epidermis of infected mice showing presence of *Leishmania* amastigotes. Scale bar: 50 μm. Circle: amastigote-containing site. Arrow: amastigotes not associated with host cells. (E) Peripheral nerve in the skin of Ctrl mice. Scale bar: 50 μm. Inset: Closer view of the nerve. Scale bar: 10 μm. (F) Peripheral nerve in the skin of challenged mice. Scale bar: 50 μm. Inset: Closer view of the nerve containing amastigotes in the vicinity. Scale bar: 25 μm. (G) Peripheral nerve in the skin of challenged mice containing an abnormal halo. Scale bar: 50 μm.

Among structures affected by the infection and inflammation, hair follicles and peripheral nerves seemed prominent, showing signs of surrounding inflammation, and deformities in sebaceous glands and arrector pili muscles (Suppl. Fig. 2A,B). Compared to Ctrl nerves (Fig. 2E), some of the nerves on the skin of infected mice showed intense surrounding inflammatory response, abnormal cellularity, seemingly bigger sizes and even presence of neighboring amastigotes (Fig. 2F). Some of them showed signs of deformity, and had an abnormal halo, indicating retraction (Fig. 2G).

Necrosis caused by *L. amazonesis* in mice is already well documented and is an important mechanism of parasite clearance during infection^30^. During our studies, some mice which had undergone the challenge with *L. amazonensis* developed necrotic lesions (Fig. 3A,D), which is apparent when compared to those that did not (Fig. 3B,E) and their contralateral paws (Fig. 3C,F). Data from mice which developed necrotic lesions was sorted from those which did not in all analyses and were compared regarding their paw withdrawal thresholds between necrotic and non-necrotic lesions, from the starting point of necrosis (Fig. 3G). No difference between thresholds was observed, showing that beyond inflammation, necrosis is also not capable of causing mechanical allodynia in *L. amazonensis* infected C57BL/6 mice. Microscopical evaluation of the skin from lesions on the infected hindpaw showed cells that could be undergoing necrosis, evidenced by swollen and fragmented nuclei, which could not be seen in Ctrl mice (Fig. 3H,I). Although intense tissue destruction is expected by the necrotic process, fibroblasts presented large nucleus and decondensed chromatin, what could be a sign of tissue remodeling, and areas of disarray, portraying a possible fibrotic process (Suppl. Fig. 2C,D).

**Fig. 3.**
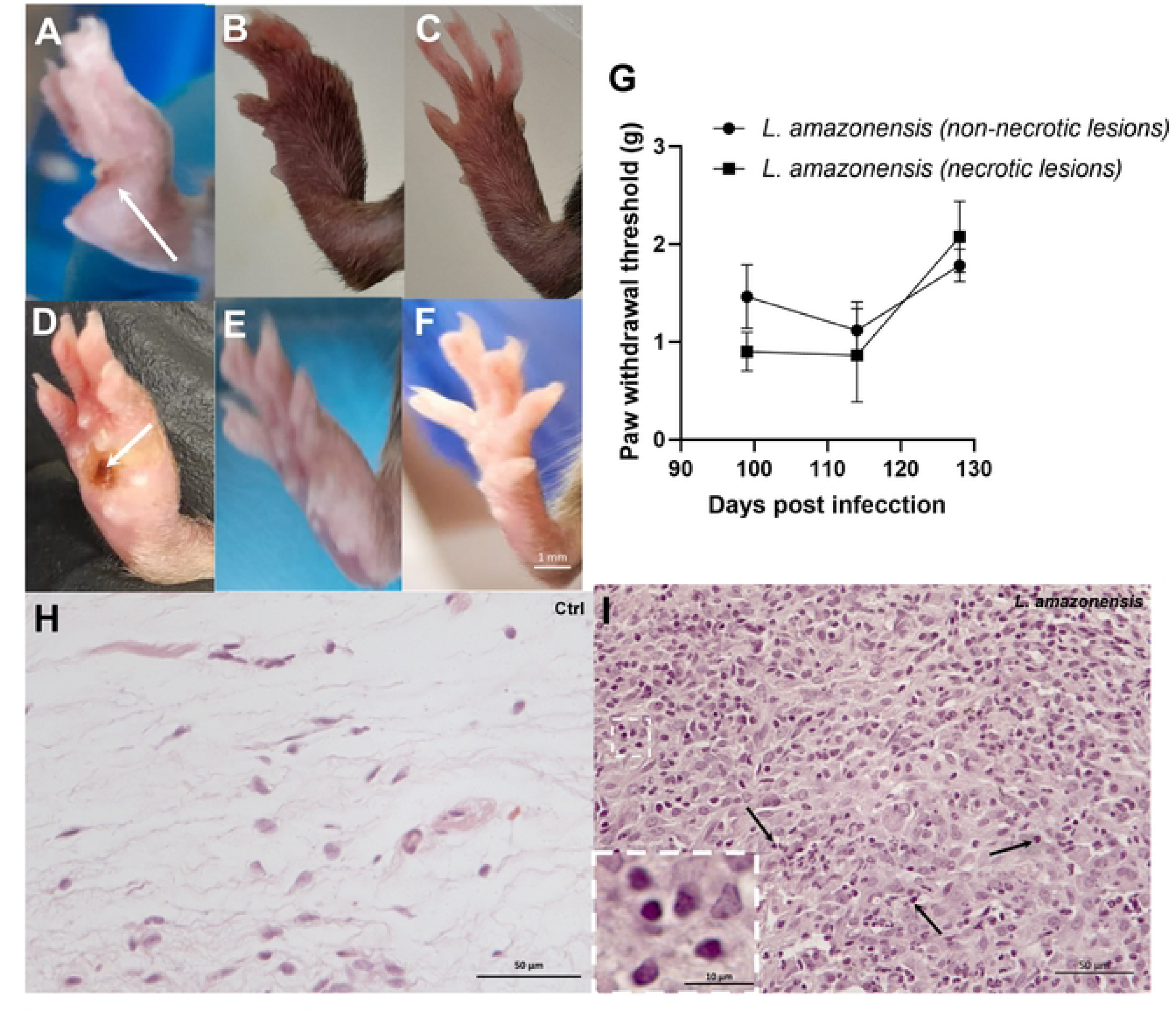
*L. amazonensis* infection causes necrosis in infected paw but does not impact on mechanical allodynia in C57BL/6 mice. (A-F) Representative images of paws from challenged mice and necrotic lesions. Scale bar = 1mm. After lesion analysis on a weekly basis, some animals presented necrotic foci in the infected paw (A,D, arrows), while others developed inflamed lesions with no signs of necrosis macroscopically (B,E). Ctrl animals are shown in (C,F). (G) The Von Frey test was conducted in both mice developing necrosis or not. First point shown in the test was the first time necrosis was seen, around 100 dpi. Data expressed by mean ± SEM, Statistical analysis through Two-way ANOVA, Sidak’s multiple comparison test, with individual variances computed for each comparison (n=3-6/group). (H-I) Representative images of H&E staining of the challenged paw after 6 months of infection. Scale bars: 50 μm. (H) Control mice. (I) C57BL/6 mice challenged with *L. amazonensis.* Cells with phenotype of being seemingly undergoing necrosis are shown.

### 3.3. *L. amazonensis* infection induces chronic phase thermal hyperalgesia dependent on IL-1β

To better evaluate the changes in nociception due to the infection, other nociceptive tests were carried out. The formalin test carried out 180 dpi showed hyperalgesia to chemical stimuli in mice challenged with *L. amazonensis* not in the first phase, characterized by direct activation of nociceptors and by the release of algogenic substances related to neuronal damage (Fig. 4A), but only in the second phase, consisting of inflammatory pain and production of inflammatory mediators (Fig. 4B). Mice also showed cold allodynia only in the chronic phase, pointed out by the acetone test (Fig. 4C, right, 120 dpi), however, cold sensitivity was not related to paw width, as shown in the correlation test (Fig. 4D).

**Fig. 4.**
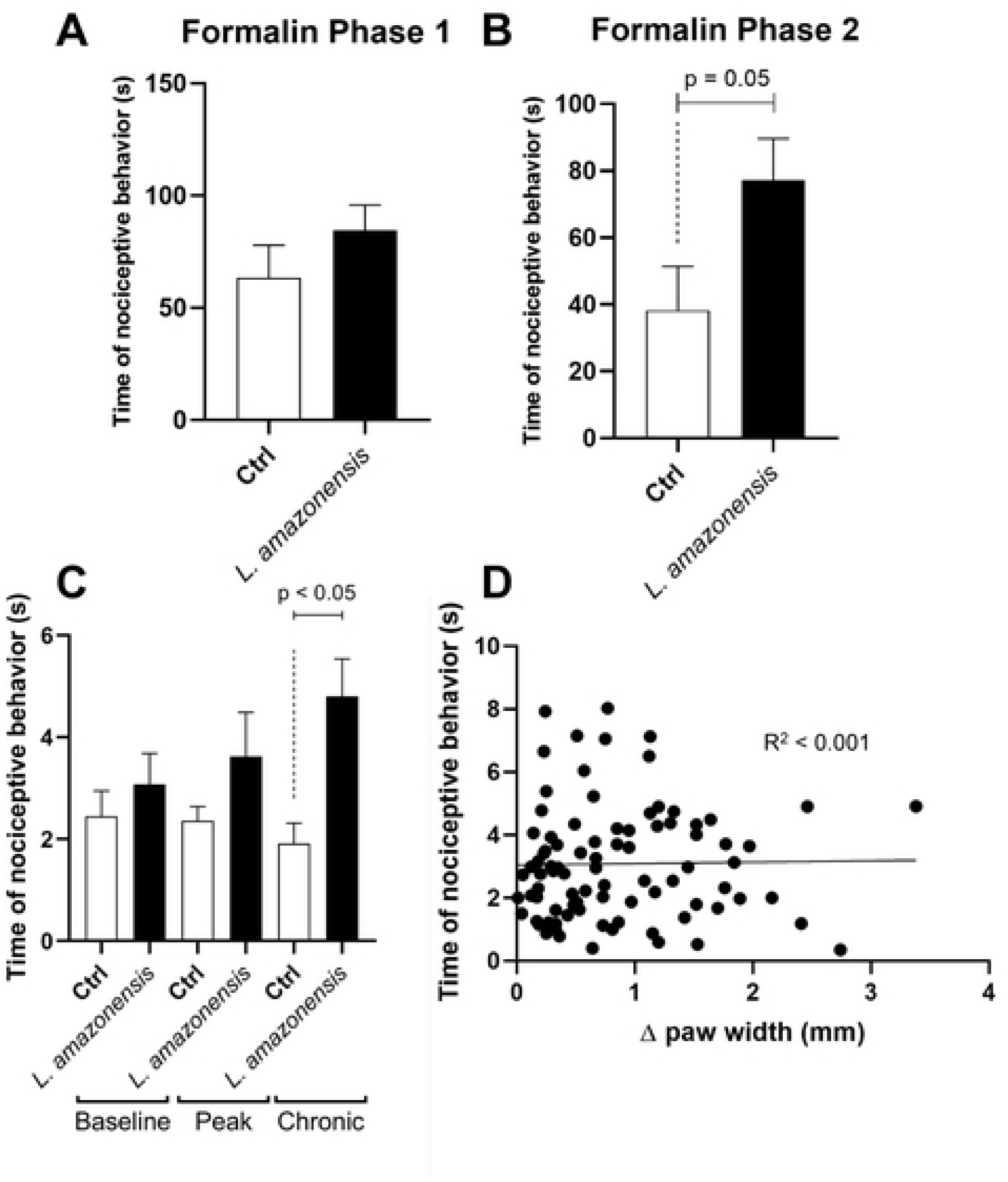
*L. amazonensis* infection leads to hypersensitization to cold and inflammatory stimuli in mice. (A,B) Time of nociceptive behavior (paw licking and shaking) in seconds after formalin injection was studied to evaluate the chemical nociception of animals in phases 1 (A), and 2 (B) of the test. Data expressed by mean ± SEM, p=0.05 (B) by Student’s t test (n=8-12/group). (C) The acetone test was performed to determine the cold nociception threshold by quantifying the reaction time in seconds. Data expressed by mean ± SEM. p=0.05 by One-way ANOVA, Tukey’s multiple comparison test, with a single pooled variance (n=7/group). (D) Correlation between time of nociceptive behavior and Δ paw width, each data point correlates to a single response in each day throughout the whole experiment, correlation tests pointed out negative results (R² < 0.001). Data expressed by individual values, and linear regression.

Surprisingly, thermal hyperalgesia in the peak phase was not evidenced by the hot plate test (Fig. 5A, left, 60 dpi). Nonetheless, the same mice showed lower thresholds regarding the thermal stimulus when evaluated at the chronic phase, (Fig. 5A, right, 120 dpi). Among the cytokines capable of leading to inflammation-induced hyperalgesia, TNF-α and IL-1β have been well characterized regarding their ability to sensitize nociceptors to upcoming noxious or innocuous stimuli^16,20^. At the chronic phase of the infection, concomitantly with thermal hyperalgesia, we found no difference in TNF-α levels in the challenged paws compared to Ctrl (Fig. 5B), but a significant increase in IL-1β levels (Fig. 5C). Regardless, paw width also showed no correlation to thermal thresholds (Fig. 5D).

**Fig. 5.**
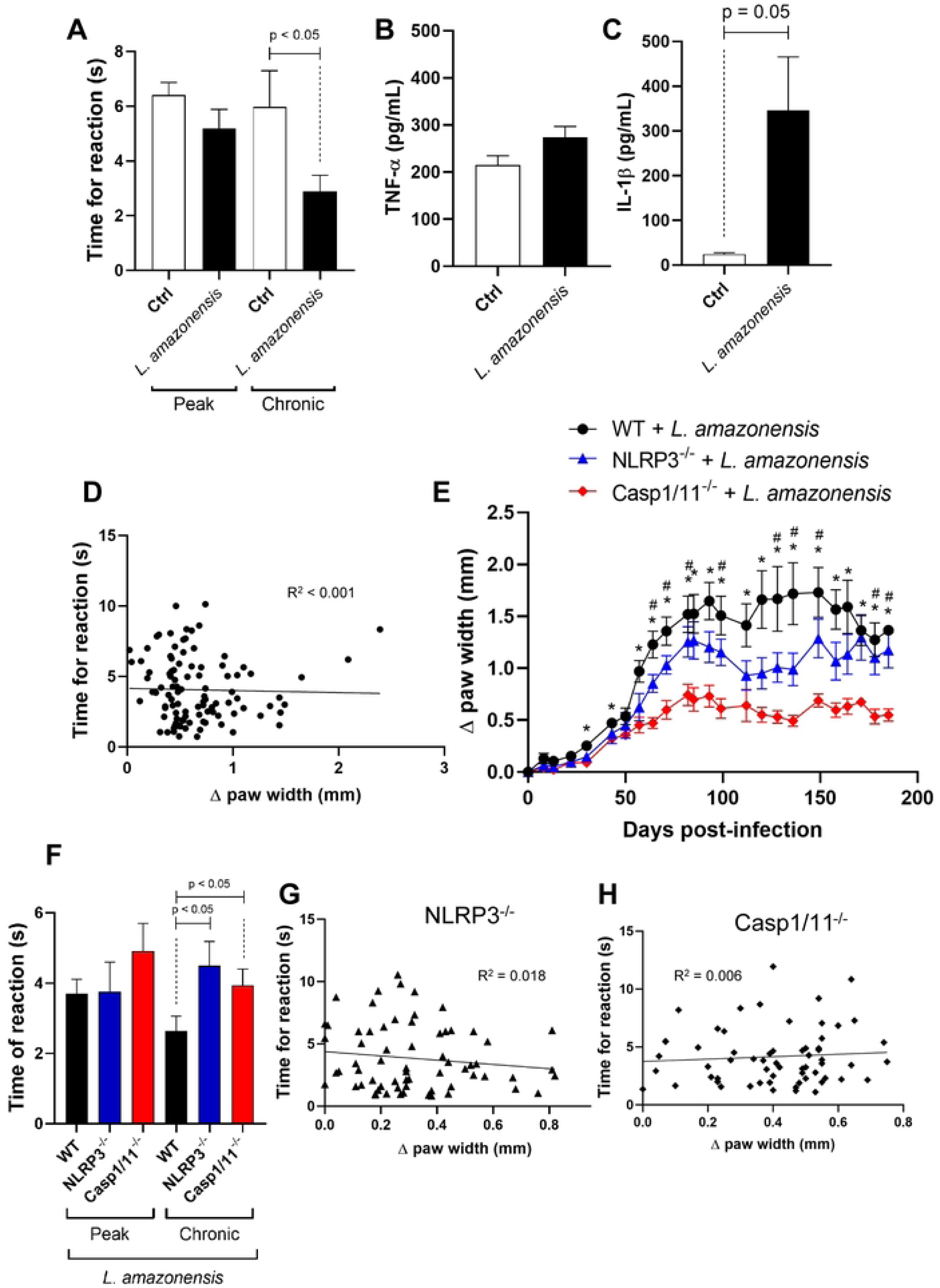
*L. amazonensis* infection induces chronic phase thermal hyperalgesia dependent on IL-1β. (A) Mice were evaluated according to their time for reaction in the hot plate test in the peak phase, 60 dpi, or chronic phase, 120 dpi. Data expressed by mean ± SEM, p<0.05 One-way ANOVA, Tukey’s multiple comparison test, with a single pooled variance (n=5-10/group). (B,C) Levels of cytokines (pg/mL) in the challenged paw were measured through ELISA of the paw homogenate 90 dpi for TNF-α (B) and IL-1β (C). Data expressed by mean ± SEM, p=0.05 (C) by Student’s t test (n=3/group). (D) Correlation between time for reaction and Δ paw width, each data point correlates to a single response in each day throughout the whole experiment, correlation tests pointed out negative results (R² < 0.001). Data expressed by individual values, and linear regression. (E) Adult WT, NLRP3^-/-^ and Casp1/11^-/-^ C57BL/6 mice were challenged subcutaneously in the right hind paw with 2 x 10^6^ cells/mL promastigotes of *Leishmania amazonensis* and were assessed throughout infection. Δ paw width was measured every week throughout the infection. Data expressed by mean ± SEM, *p<0.05 (WT versus Casp1/11^-/-^), #p<0.05 (NLRP3^-/-^ versus Casp1/11^-/-^) by Two-way ANOVA, Sidak’s multiple comparison test, with individual variances computed for each comparison (n=7-11/group). Additionally, by Tukey’s multiple comparison (main column effect), NLRP3^-/-^ mice presented overall lower Δ paw width than WT (p<0.01) and higher than Casp1/11^-/-^ (p<0.0001). (F) Mice were evaluated according to their time for reaction in the hot plate test in the peak phase, 60 dpi, or chronic phase, 120 dpi. Data expressed by mean ± SEM, p<0.05 One-way ANOVA, Tukey’s multiple comparison test, with a single pooled variance (n=7-15/group). (G,H) Correlation between time for reaction and Δ paw width for NLRP3^-/-^ (G) and Casp1/11^-/-^ (H), each data point correlates to a single response in each day throughout the whole experiment, correlation tests pointed out negative results (R²=0.018 and R²=0.006, respectively). Data expressed by individual values, and linear regression.

For better assaying IL-1β role in the chronic phase thermal hyperalgesia caused by *L. amazonensis,* we used different knockout (KO) mice lacking the production of IL-1β due to deficiency in the correct activation and assembly of the inflammasome, NLRP3^-/-^ and the double KO Casp1/11^-/-^ mice. Paw width due to infection had lower overall growth in Casp1/11^-/-^ mice compared to WT, but this was also true when compared to NLRP3^-/-^ mice (Fig. 5E). NLRP3^-/-^ mice also showed lower paw width growth compared to WT mice, however at lower levels than the difference seen for WT vs Casp1/11^-/-^. The reduced inflammation observed was also present in microscopical analysis, despite hair follicles and nerve terminals also appearing to be committed (Suppl. Fig. 2D-F). Thresholds for the heat stimulus were at equal levels in the peak phase comparing WT and KO mice (Fig. 5F, left, 60 dpi), however thresholds of WT mice were lower than both NLRP3^-/-^ and Casp1/11^-/-^ mice in the chronic phase (Fig. 5F, right, 120 dpi), suggesting the thermal hyperalgesia caused by *L. amazonensis* infection was reduced when there was deficiency in the production of IL-1β. The correlation between KO mice regarding thermal thresholds and paw width was negative (Fig. 5G,H).

This thermal hyperalgesia is probably directly associated with the IL-1β, since parasite load among these mice did not differ, showing this effect had no correlation with the presence of parasites, either in the challenged paws or the draining lymph nodes of infected mice (Suppl. Fig. 3)

Interestingly, neither mechanical nor cold sensitivity were affected by the KOs, regardless of the phase (Fig. 6A and 6D, respectively). Seemingly, KO mice also did not present correlation between paw width and mechanical thresholds (Fig. 6B,C), however NLRP3^-/-^ mice showed a trend for increased sensitivity to cold when paw width was higher (Fig. 6E, R²=0.23), what was not observable for Casp1/11^-/-^ mice (Fig. 6F).

**Fig. 6.**
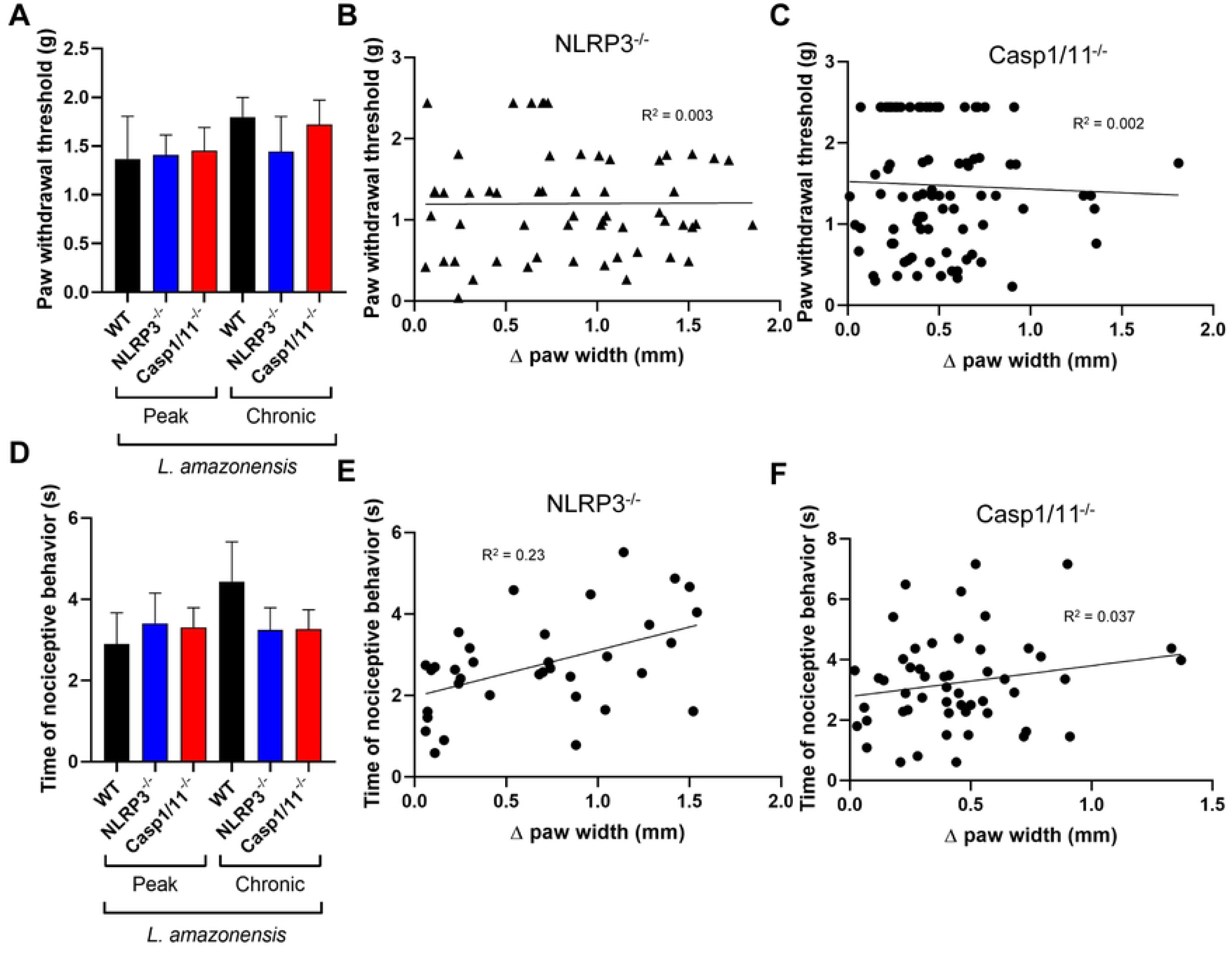
Deficiency in IL-1β production does not affect other forms of sensitivity. (A-F) WT, NLRP3^-/-^ and Casp1/11^-/-^ mice were evaluated regarding other forms of nociception. (A) Response threshold to the Von Frey test in the peak phase, 60 dpi, or chronic phase, 120 dpi. Data expressed by mean ± SEM, One-way ANOVA, Tukey’s multiple comparison test, with a single pooled variance (n=5-11/group). (B,C) Correlation between paw withdrawal threshold upon mechanical stimulus and Δ paw width for NLRP3^-/-^ (B) and Casp1/11^-/-^ (C), each data point correlates to a single response in each day throughout the whole experiment, correlation tests pointed out negative results (R²=0.003 and R²=0.002, respectively). Data expressed by individual values, and linear regression. (D) Nociceptive behavior in seconds to the Von Frey test in the peak phase, 60 dpi, or chronic phase, 120 dpi. Data expressed by mean ± SEM, One-way ANOVA, Tukey’s multiple comparison test, with a single pooled variance (n=5-11/group). (E,F) Correlation between Nociceptive behaviorNociceptive behavior upon cold stimulus and Δ paw width for NLRP3^-/-^ (E) and Casp1/11^-/-^ (F), each data point correlates to a single response in each day throughout the whole experiment, correlation tests pointed out weak correlation or negative results (R²=0.23, p<0.01, and R²=0.037, respectively). Data expressed by individual values, and linear regression.

### 3.4. Selective thermal hyperalgesia in the chronic phase of *L. amazonensis* infection in C57BL/6 mice is caused by direct activation of IL-1RI^+^ thermosensitive nociceptors in the dorsal root ganglia

We have successfully proven that *L. amazonensis* infection in C57BL/6 mice does not induce mechanical allodynia, however, it leads to thermal hyperalgesia exclusively in its chronic phase. Nonetheless, how both these phenomena can happen simultaneously is still not fully understood. To confirm the mechanism of chronic phase thermal hyperalgesia, we conducted the hot plate test in different KO animals participating in the production or recognition of IL-1β and IL-18 downstream after Caspase-1 activation, IL-1R^-/-^, IL-18^-/-^ and IL-18R^-/-^. IL-1R is the IL-1β receptor, and when activated, it can reduce activation thresholds or even directly activating nociceptors^31,32^. IL-18 is a cytokine other than IL-1ß activated by the inflammasome pathway, and its mechanism of action in inflammation relies on its interaction with its receptor IL-18R. Despite paw Δ paw widths did not differ among groups (Fig. 7A), both IL-18^-/-^ and IL-18R^-/-^ mice showed no changes in their thermal thresholds when compared to WT at 120 dpi (Fig. 7B). Strikingly, IL-1R^-/-^ mice showed a great increase to its threshold, meaning its absence was sufficient to alleviate chronic phase thermal hyperalgesia, which was seemingly not associated with inflammation. Since inflammation of the lesion site is not a defining factor for chronic phase hyperalgesia, we hypothesized IL-1β was acting in other regions involved in the nociceptive pathway. The dorsal root ganglion (DRG) is located next to the spinal cord, and as a converging point in the afferent pathway, it is composed of multiple cell bodies of nociceptors, many of which innervate the skin^33^. Immunofluorescence for IL-1R^+^ TRPV1^+^ neurons in the spinal cord revealed their presence in both WT and Casp1/11^-/-^ mice, albeit seemingly more evident in the latter (Suppl. Fig. 4). We then evaluated whether there were changes in the DRG and adjacencies by qPCR analysis of expression levels in WT and Casp1/11^-/-^ mice, Ctrl or challenged with *L. amazonensis*. Levels of IL-1R were not altered among WT Ctrl or challenged mice, however in Casp1/11^-/-^ mice, infection led to increased expression levels of this gene in most individuals, although not statistically significant (Fig. 7C). Regarding IL-1β expression, most WT challenged mice also showed an increase, although not statistically, whilst Casp1/11^-/-^ mice showed minimal levels of expression of this cytokine (Fig. 7D). Finally, since the transient receptor potential vanilloid 1 (TRPV1) has been previously shown to be associated to IL-1R in IL-1β-responsible neurons in the DRG during chronic inflammation^34^, we also assayed its expression levels in the DRGs. This ion channel was approximately 3-fold upregulated in WT mice challenged with *L. amazonensis* compared to Ctrl (Fig. 7E), pointing to a sensitization-prone environment for thermosensitive neurons.

**Fig. 7.**
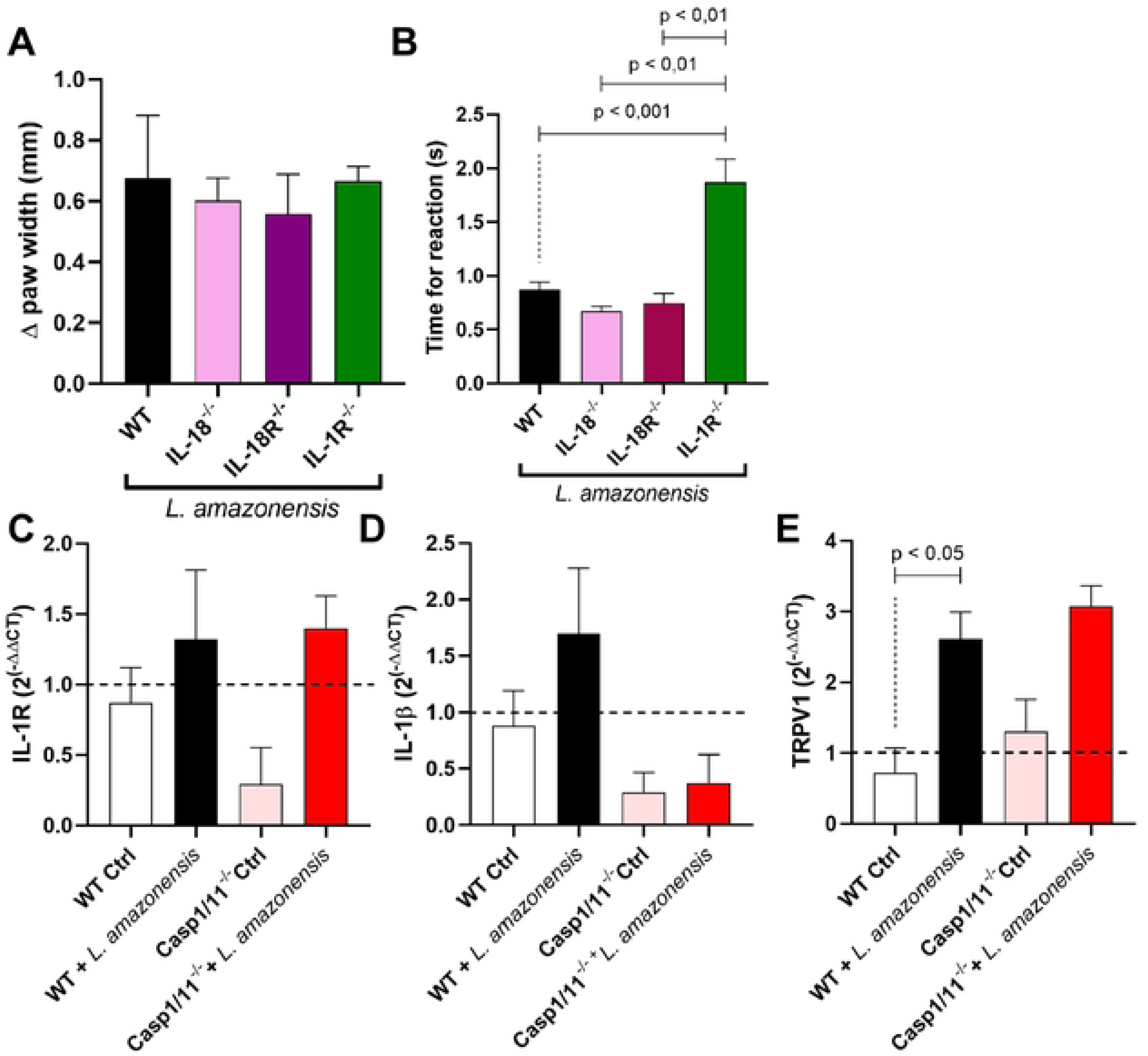
Selective thermal hyperalgesia in the chronic phase of *L. amazonensis* infection in C57BL/6 mice is caused by direct activation of IL-1RI^+^ thermosensitive nociceptors in the DRGs. (A,B) Adult WT, IL-18^-/-^, IL-18R^-/-^ and IL-1R^-/-^ C57BL/6 mice were challenged subcutaneously in the right hind paw with 2 x 10^6^ cells/mL promastigotes of *Leishmania amazonensis* and were assessed regarding time for Δ paw width at 120 dpi (A), or reaction to the hot plate test (B). Data expressed by mean ± SEM, p<0.01 and p<0.001 One-way ANOVA, Tukey’s multiple comparison test, with a single pooled variance (n=5-15/group). (C-E) Fold change of the IL-1R (C), IL-1β (D) and TRPV1 (E) encoding genes was calculated by 2^(-ΔΔCT)^, and normalized from the Ctrl group. Mice analyzed were either WT or Casp1/11^-/-^, Ctrl or challenged with *L. amazonensis*. Data expressed by mean ± SEM, p<0.05 One-way ANOVA, Tukey’s multiple comparison test, with a single pooled variance (n=3-5/group).

## 4. Discussion

Patients suffering from LCL usually develop painless ulcerated nodules, despite intense inflammation and necrosis associated with these lesions^35^. The mechanisms underlying the absence of mechanical allodynia are still unclear. Most experimental studies in non-human laboratory animals are based on BALB/c mice, which harbors a susceptible environment for the infection with *Leishmania* parasites, pointing to infection-induced hyperalgesia^14,15^. However, C57BL/6 mice respond differently to the infection with *L. amazonensis*, with a partially resistant phenotype^36,37^, being closely related to the response observed in patients ^38^.

Through the Von Frey test, we evaluated the mechanical response thresholds of *L. amazonensis* challenged mice upon innocuous mechanical stimulus throughout the infection. Mechanical allodynia was never present in challenged mice compared to Ctrl and together with that, mice showed no alteration in their tactile behavior. The absence of correlation between paw width, necrosis, and mechanical thresholds further reinforces that usual pain-inducing pathways in the lesion site are not sufficient to induce hypersensitization in *L. amazonensis* experimental infection. Furthermore, recent studies suggest that *Leishmania* infection induces pain-reducing reprogramming by metabolic studies^9^. Microscopical evaluation of tissue in the lesion site showed morphological alterations that may enlighten the mechanisms underlying these findings, namely evidence for fibroplasia and apparent damage to hair follicles and nerve terminals. Hair follicles play an important role in touch sensitivity, as well as nerve terminals such as the Ruffini, Pacini and Meissner corpuscles, and free nerve endings, all of which if impaired, may also dampen the ability of hypersensitization of nerves contained in these structures. Coupled with degeneration, tissue remodeling and activation of fibroblasts may lead to replacement of previous functional structures for non-specialized conjunctive tissue, dampening further the ability to respond to hypersensitization signals^39,40^.

Sensitivity loss due to nerve changes in skin is a phenomenon also observed for another important infectious disease, Hansen’s disease. Patients committed by this disease may develop ulcerating cutaneous lesions that are impervious to pain^3^. This absence of pain in the ulcers of these patients is linked to neuropathies and changes in the nerves and nerve endings^41^.

Even though we found evidence for desensitization in the lesion site to noxious stimuli, thermal hyperalgesia is ubiquitous in studies with mice^16,21,42^. Even in patients, treatment with thermotherapy, despite proven efficient, is acknowledged as painful^43^. Therefore, absence of mechanical allodynia is not the only nociceptive change induced by *L. amazonensis* infection, and thus another mechanism leads to selective hyperalgesia.

In fact, we found no evidence of thermal hyperalgesia, neither cold nor heat, in the phase when lesion size peaked, around 60 dpi. However, starting at the chronic phase, around 120 dpi, thermal hyperalgesia both to cold and heat stimuli was present, concomitantly with higher levels of IL-1β. Mice also showed no signs of increased sensitivity to immediate formaldehyde sensing, but instead, were more sensitive to formaldehyde-induced production of inflammatory mediators. Mice deficient in the IL-1β production pathway showed no infection-induced hyperalgesia, further reinforcing its importance in chronic phase hyperalgesia. Importantly, these mice also showed alterations in the morphology of lesions, and thus we expect them to harbor the sensitization-resistant phenotype in the lesion site. The fact that parasite levels remained the same in the paws and lymph nodes of chronically infected mice among WT and KO further reinforces the direct effects of IL-1β-driven selective thermal hyperalgesia.

IL-1β is capable of directly activating nociceptors and lowering their activation thresholds ^31,34^, but also of initiating the TNF receptor-associated factor 6 (TRAF6) signaling cascade, culminating in the activation of proinflammatory transcription factors (32). Surprisingly, TNF-α levels were not increased. This cytokine can induce hypersensitization of nociceptors through three main mechanisms: direct binding to its receptors in neurons, leading to reduction in the activation thresholds of NaV1.8^44^; induction of inflammation through TNFR1 and TNFR2 signaling cascade, culminating in activation of proinflammatory transcription factors^45^; and inducing the production of IL-1β^46^. Since TNF-α was not upregulated, these data provide evidence that inflammation-induced IL-1β-dependent hyperalgesia during *L. amazonensis* infection depends on its direct action through receptor binding.

IL-1β receptors, the IL-1Rs, present two forms: IL-1RI, expressed in T lymphocytes, fibroblasts, epithelial cells, endothelial cells, and neurons, and IL-1RII, expressed in B lymphocytes and APCs^47^. In order to confirm whether direct activation of IL-1Rs led to thermal hyperalgesia at the chronic phase, we used mice KO for IL-1R, and to further prove the involvement in this process is solely of IL-1β, and not pyroptosis or inflammation, we used KO mice for IL-18, the other cytokine activated by the inflammasome, and its receptor, IL-18R. Indeed, IL-1R deficiency increased thermal thresholds compared to WT, IL-18 and IL-18R, significantly dampening infection-induced hyperalgesia.

A specific population of TRPV1^+^ IL-1R^+^ nociceptors has been experimentally characterized previously in chronic inflammatory diseases studies in mice^34,48^. This population was present in the DRGs and presented close association to microglia, as well as being responsive to IL-1β. Interestingly, TRPV1 is also a heat-responsive ion channel, and thus nociceptors expressing this protein are expected to respond to noxious heat. Nonetheless, *L. amazonensis* infection was shown to induce thermal hyperalgesia through IL-1β production in the DRGs^14^. When evaluating gene expression in the DRGs in the chronic phase, we found TRPV1 was upregulated. IL-1β and IL-1R expression were not statistically increased, but most infected mice showed higher expression levels when compared to their Ctrl counterparts.

Some C-type fibers are unresponsive to mechanical stimulus, but responsive to thermal^49^. We found evidence that IL-1β is capable of selectively inducing thermal hyperalgesia. Interestingly, Hansen’s disease is also associated with chronic pain.

With these findings, we hypothesize for the first time that *L. amazonensis* infection leads to both desensitization and hypersensitization, but through different mechanisms. The first mechanism of nociceptive change in *L. amazonensis* infection relies on general desensitization through changes in the skin in the lesion size, affecting nerve terminals responsible for pain signaling. The second mechanism is IL-1β dependent selective thermal hyperalgesia, through sensitization of IL-1R-bearing neurons in the DRGs, which may be mechano-insensitive.

## Limitations of the study

We acknowledge the limitation imposed by sex restriction upon the choice of only females in our studies. Indeed, sex is a factor that can influence pain, however other factors must be considered and may have an even greater impact ^50^.

One major limitation was the use of the contralateral paw as control in many of the data presented to increase the sample size. Indeed, it is important to note that systemic inflammation due to chronic infection is expected, possibly affecting contralateral paws, which were used as control in many of the experiments. However, parasite load could not be observed in the contralateral paws of chronically infected mice (Suppl. Fig. 3). Nonetheless, we also compared previously the ipsilateral paws of mice challenged with PBS mice, and the contralateral paws of *L. a.* challenged mice regarding mechanical and cold allodynia (Suppl. Fig. 5), and since they presented no statistical differences regarding their response, we are secure that using contralateral paws in nociceptive studies using *L. amazonensis* infected C57BL/6 mice is a valid form of control.

Lastly, histological and immunohistochemical results could not be quantified due to internal limitations. We acknowledge that it would be greatly beneficial to our work for confirming our observations analyzing fibrosis, necrosis, nerve staining and DRG immunostaining. Nevertheless, it was possible to give a wide view of changes in the tissue that allied to our nociceptive data helped us understand the general process underlying pain in *L. amazonensis* infection, despite more data and conclusive studies are needed to confirm our hypothesis.

## Acknowledgements

We thank the financial support from Brazilian funding agencies: Fundação de Amparo à Pesquisa do Estado do Rio de Janeiro (FAPERJ E-26/010.101087/2018, Brazil), Conselho Nacional de Desenvolvimento Científico e Tecnológico (CNPq 308012/2019-4, Brazil) and Fundação Coordenação de Aperfeiçoamento de Pessoal de Nível Superior (CAPES, Brazil) Finance code 001. We also thank the Histotechnology Platform (Fiocruz, Pavilion Cardoso Fontes, RJ) for the processing and staining of skin tissue.

---

**Suppl. Fig. 1. Experimental design.** Adult WT, IL-18^-/-^, IL-18R^-/-^ and IL-1R^-/-^ C57BL/6 mice were challenged subcutaneously in the right hind paw with 2 x 10^6^ cells/mL promastigotes of *Leishmania amazonensis*, or PBS as control. Behavioral assessments were conducted through the infection (Von Frey test, acetone test, hot plate test and formalin test) until mice were infected for 180 days, when they were euthanized. After euthanasia, tissues were collected and used for IHC, qPCR and ELISA tests.

**Suppl. Fig. 2. Hair follicles were compromised and fibroblasts presented an active phenotype in later phases of *Leishmania amazonensis* infection.** Adult C57BL/6 mice were challenged subcutaneously in the right hind paw with 2 x 10^6^ cells/mL promastigotes of *Leishmania amazonensis* and tissue was collected 6 months after challenge. Representative images of H&E staining, with scale bars represented in the lower right. (A,B) Skin tissue of Ctrl (A) or challenged (B) mice. Scale bar: 100 μm (A) or 50 μm (B). Arrows: points to a hair follicle. (C,D) Images of active fibroblasts in the skin of WT (C) and Casp1/11^-/-^ mice (D). Scale bar: 50 μm. Insets: closer view of fibroblasts undergoing fibroplasia. Scale bar: 25 μm. (E,F) Peripheral nerve of Ctrl (E) and Casp1/11^-/-^ mice challenged with *L. amazonensis* (F). Scale bar: 50 μm. Arrows: peripheral nerves. Inset: Closer view of peripheral nerves. Scale bar: 10 μm (E), 25 μm (F).

**Suppl. Fig. 3. Parasite load did not differ in paws and lymph nodes among chronically infected mice.** Parasite load was determined by LDA of chronically infected mice (180 dpi) at the paws (A,C) and lymph nodes (B,D), in WT, NLRP3^-/-^ and Casp1/11^-/-^ mice (A,B) and WT, IL-18^-/-^, IL-18R^-/-^ and IL-1R^-/-^ mice (C,D). Data expressed by mean ± SEM. One-way ANOVA, Tukey’s multiple comparison test, with a single pooled variance (n=3-5/group).

**Suppl. Fig. 4. Presence of IL-1R1^+^ TRPV1^+^ neurons in the spinal cord of L. amazonensis challenged mice in the chronic phase of the infection.** (A,B) Immunofluorescence confocal microscopy of spinal cord sections from WT or Casp1/11^-/-^ C57BL/6 mice challenged with 2 x 10^6^ cells/mL promastigotes of *Leishmania amazonensis* after 6 months of infection. TRPV1 (green) and IL-1R1 (red) were colocalized in neurons in both mice, despite an apparent higher intensity in WT compared to Casp1/11^-/-^ mice. Scale bar: 20µm.

**Suppl. Fig. 5. Paws from mice challenged with PBS presented similar response compared to contralateral paws from mice challenged with *L. amazonensis*.** Mice were either challenged with 20µL PBS or 2 x 10^6^ cells/mL promastigotes of *Leishmania amazonensis* and compared between the ipsilateral paw or contralateral paw, respectively. Analyses were conducted in the early phase (10 dpi) or chronic phase (120 dpi) for both mechanical (A) or cold (B) allodynia. Data expressed by mean ± SEM. One-way ANOVA, Tukey’s multiple comparison test, with a single pooled variance (n=5-10/group).

## Notes

### Competing Interest Statement

The authors have declared no competing interest.

## References

1. Bezerra JMT, de Araújo VEM, Barbosa DS, Martins-Melo FR, Werneck GL, Carneiro M. Burden of leishmaniasis in Brazil and federated units, 1990-2016: Findings from Global Burden of Disease Study 2016. PLoS Negl Trop Dis. 2018;12(9):e0006697-. 10.1371/journal.pntd.0006697

2. Organization PAH, Organization PAH. Leishmaniasis: Epidemiological Report for the Americas. No.12 (December 2023).https://iris.paho.org/handle/10665.2/59155. 2023.

3. Torres-Guerrero E, Quintanilla-Cedillo MR, Ruiz-Esmenjaud J, Arenas R. Leishmaniasis: a review. F1000Res. 2017;6(May):750. doi:10.12688/f1000research.11120.1

4. Hashim FA, Ahmed AE, El Hassan M, et al. Neurologic changes in visceral leishmaniasis. American Journal of Tropical Medicine and Hygiene. 1995;52(2):149–154. doi:10.4269/ajtmh.1995.52.149

5. Petersen CA, Greenlee MHW. Neurologic Manifestations of Leishmania spp. Infection. J Neuroparasitology. 2011;2(Vl):1–5. doi:10.4303/jnp/n110401

6. de Aragão REM, Barreira IMA, Pereira LA, et al. Intraretinal hemorrhage associated with visceral leishmaniasis. Rev Bras Oftalmol. 2015;74(6):393–395. doi:10.5935/0034-7280.20150083

7. Kubba R, El-Hassan AM, Al-Gindan Y, Omer AHS, Bushra M, Kutty MK. Peripheral Nerve Involvement in Cutaneous Leishmaniasis (Old World). Int J Dermatol. 1987;26(8):527–531. doi:10.1111/j.1365-4362.1987.tb02295.x

8. Bussmann AJC, Santos LFS, Ferreira RN, et al. Leishmania spp. amastigotes surrounding sensory nerve fibers in human painless skin ulcers: Evidence of pathogen-neuron proximity and absence of neuronal apoptosis. Acta Trop. 2024;256. doi:10.1016/j.actatropica.2024.107265

9. Volpedo G, Oljuskin T, Cox B, et al. Leishmania mexicana promotes pain-reducing metabolomic reprogramming in cutaneous lesions. iScience. 2023;26(12). doi:10.1016/j.isci.2023.108502

10. Satti MB, El-Hassan AM, Al-Gindan Y, Osman MA, Al-Sohaibani MO. Peripheral Neural Involvement in Cutaneous Leishmaniasis. Int J Dermatol. 1989;28(4):243–247. doi:10.1111/j.1365-4362.1989.tb04813.x

11. Abreu-Silva AL, Calabrese KS, Tedesco RC, Mortara RA, Gonçalves Da Costa SC. Central nervous system involvement in experimental infection with Leishmania (Leishmania) amazonensis. American Journal of Tropical Medicine and Hygiene. 2003;68(6):661–665. doi:10.4269/ajtmh.2003.68.661

12. Borghi SM, Fattori V, Conchon-Costa I, Pinge-Filho P, Pavanelli WR, Verri WA. Leishmania infection: painful or painless? Parasitol Res. 2017;116(2):465–475. doi:10.1007/s00436-016-5340-7

13. Cangussú SD, Souza CC, Castro MSA, et al. The endogenous cytokine profile and nerve fibre density in mouse ear Leishmania major-induced lesions related to nociceptive thresholds. Exp Parasitol. 2013;133(2):193–200. doi:10.1016/j.exppara.2012.11.015

14. Borghi SM, Fattori V, Pinho-Ribeiro FA, et al. Contribution of spinal cord glial cells to L. amazonensis experimental infection-induced pain in BALB/c mice. J Neuroinflammation. 2019;16(1):1–23. doi:10.1186/s12974-019-1496-2

15. Karam MC, Hamdan HG, Abi Chedid NA, Bodman-Smith KB, Eales-Reynolds LJE, Baroody GM. Leishmania major: Low infection dose causes short-lived hyperalgesia and cytokines upregulation in mice. Exp Parasitol. 2006;113(3):168–173. doi:10.1016/j.exppara.2006.01.003

16. Borghi SM, Fattori V, Ruiz-Miyazawa KW, et al. Leishmania (L). amazonensis induces hyperalgesia in balb/c mice: Contribution of endogenous spinal cord TNFα and NFκB activation. Chem Biol Interact. 2017;268:1–12. doi:10.1016/j.cbi.2017.02.009

17. Leung S, Liu X, Fang L, Chen X, Guo T, Zhang J. The cytokine milieu in the interplay of pathogenic Th1/Th17 cells and regulatory T cells in autoimmune disease. Cell Mol Immunol. 2010;7(3):182–189. doi:10.1038/cmi.2010.22

18. Tomiotto-Pellissier F, Bortoleti BT da S, Assolini JP, et al. Macrophage Polarization in Leishmaniasis: Broadening Horizons. Front Immunol. 2018;9. doi:10.3389/fimmu.2018.02529

19. Zamboni DS, Sacks DL. Inflammasomes and Leishmania: in good times or bad, in sickness or in health. Curr Opin Microbiol. 2019;52:70–76. doi:10.1016/j.mib.2019.05.005

20. Cunha TM, Verri WA, Silva JS, Poole S, Cunha FQ, Ferreira SH. A cascade of cytokines mediates mechanical inflammatory hypernociception in mice. Proc Natl Acad Sci U S A. 2005;102(5):1755–1760. doi:10.1073/pnas.0409225102

21. Kanaan SA, Saadé NE, Karam M, Khansa H, Jabbur SJ, Jurjus AR. Hyperalgesia and upregulation of cytokines and nerve growth factor by cutaneous leishmaniasis in mice. Pain. 2000;85(3):477–482. doi:10.1016/S0304-3959(99)00297-3

22. Broz P, Dixit VM. Inflammasomes: Mechanism of assembly, regulation and signalling. Nat Rev Immunol. 2016;16(7):407–420. doi:10.1038/nri.2016.58

23. Maia JRLCB, Machado LKA, Fernandes GG, et al. Mitotherapy prevents peripheral neuropathy induced by oxaliplatin in mice. Neuropharmacology. 2024;245:109828. 10.1016/j.neuropharm.2023.109828

24. Chaplan SR, Bach FW, Pogrel JW, Chung JM, Yaksh TL. Quantitative assessment of tactile allodynia in the rat paw. J Neurosci Methods. 1994;53(1):55–63. doi:10.1016/0165-0270(94)90144-9

25. McNamara CR, Mandel-Brehm J, Bautista DM, et al. TRPA1 mediates formalin-induced pain. Proceedings of the National Academy of Sciences. 2007;104(33):13525–13530. doi:10.1073/pnas.0705924104

26. Tjølsen A, Berge OG, Hunskaar S, Rosland JH, Hole K. The formalin test: an evaluation of the method. Pain. 1992;51(1):5–17. doi:10.1016/0304-3959(92)90003-T

27. Dias FC, Alves VS, Matias DO, et al. The selective TRPV4 channel antagonist HC-067047 attenuates mechanical allodynia in diabetic mice. Eur J Pharmacol. 2019;856:172408. 10.1016/j.ejphar.2019.172408

28. Fontes-Dantas FL, Fernandes GG, Gutman EG, et al. SARS-CoV-2 Spike protein induces TLR4-mediated long-term cognitive dysfunction recapitulating post-COVID-19 syndrome in mice. Cell Rep. 2023;42(3). doi:10.1016/j.celrep.2023.112189

29. Punchard NA, Whelan CJ, Adcock I. The Journal of Inflammation. J Inflamm. 2004;1. doi:10.1186/1476-9255-1-1

30. Andrade ZA, Reed SG, Roters B. PATOGENIA DA LEISHMANIOSE CUTÂNEA EXPERIMENTAL. A IMPORTÂNCIA DA NECROSE NA ELIMINAÇÃO DOS PARASITOS DAS LESÕES. Vol 17.; 1984.

31. Obreja O, Rathee PK, Lips KS, Distler C, Kress M. IL-1J potentiates heat-activated currents in rat sensory neurons: involvement of IL-1RI, tyrosine kinase, and protein kinase C. The FASEB Journal. 2002;16(12):1497–1503. 10.1096/fj.02-0101com

32. Sakurai H. Targeting of TAK1 in inflammatory disorders and cancer. Trends Pharmacol Sci. 2012;33(10):522–530. doi:10.1016/j.tips.2012.06.007

33. Haberberger RV, Barry C, Dominguez N, Matusica D. Human dorsal root ganglia. Front Cell Neurosci. 2019;13. doi:10.3389/fncel.2019.00271

34. Mailhot B, Christin M, Tessandier N, et al. Neuronal interleukin-1 receptors mediate pain in chronic inflammatory diseases. Journal of Experimental Medicine. 2020;217(9):1–19. doi:10.1084/jem.20191430

35. Peninsula Y, Fernando M, Andrade-Narváez J, Vargas-González A, Canto-Lara SB, Damián-Centeno AG. Clinical Picture of Cutaneous Leishmaniases Due to Leishmania (Leishmania) Mexicana in The. Vol 96.; 2001.

36. Sacks D, Noben-Trauth N. The immunology of susceptibility and resistance to Leishmania major in mice. Nat Rev Immunol. 2002;2(11):845–858. doi:10.1038/nri933

37. VELASQUEZ LG, GALUPPO MK, DE REZENDE E, et al. Distinct courses of infection with Leishmania (L.) amazonensis are observed in BALB/c, BALB/c nude and C57BL/6 mice. Parasitology. 2016;143(6):692-703. doi:DOI: 10.1017/S003118201600024X

38. Alexander J, Brombacher F. T helper1/T helper2 cells and resistance/susceptibility to Leishmania infection: Is this paradigm still relevant? Front Immunol. 2012;3(APR). doi:10.3389/fimmu.2012.00080

39. Themistocleous AC, Ramirez JD, Shillo PR, et al. The Pain in Neuropathy Study (PiNS): A cross-sectional observational study determining the somatosensory phenotype of painful and painless diabetic neuropathy. Pain. 2016;157(5):1132–1145. doi:10.1097/j.pain.0000000000000491

40. Hsieh ST, Chiang HY, Lin WM. *Pathology of Nerve Terminal Degeneration in the Skin*. Vol 59.; 2000. https://academic.oup.com/jnen/article/59/4/297/2609881

41. Gurel MS, Tekin B, Uzun S. Cutaneous leishmaniasis: A great imitator. Clin Dermatol. 2020;38(2):140–151. 10.1016/j.clindermatol.2019.10.008

42. Karam MC, Merckbawi R, Salman S, Mobasheri A. Atenolol reduces Leishmania major-induced hyperalgesia and TNF-α without affecting IL-1β or keratinocyte derived chemokines (KC). Front Pharmacol. 2016;7(FEB):1–10. doi:10.3389/fphar.2016.00022

43. Gonçalves SVCB, Costa CHN. Treatment of cutaneous leishmaniasis with thermotherapy in Brazil: An efficacy and safety study. An Bras Dermatol. 2018;93(3):347–355. doi:10.1590/abd1806-4841.20186415

44. Oliveira Júnior JO de, Portella Junior CSA, Cohen CP. Inflammatory mediators of neuropathic pain. Revista Dor. 2016;17. doi:10.5935/1806-0013.20160045

45. Jang DI, Lee AH, Shin HY, et al. The role of tumor necrosis factor alpha (Tnf-α) in autoimmune disease and current tnf-α inhibitors in therapeutics. Int J Mol Sci. 2021;22(5):1–16. doi:10.3390/ijms22052719

46. Cunha FQ, Poole S, Lorenzetti BB, Ferreira SH. The pivotal role of tumour necrosis factor α in the development of inflammatory hyperalgesia. Br J Pharmacol. 1992;107(3):660–664. doi:10.1111/j.1476-5381.1992.tb14503.x

47. Kaneko N, Kurata M, Yamamoto T, Morikawa S, Masumoto J. The role of interleukin-1 in general pathology. Inflamm Regen. 2019;39(1). doi:10.1186/s41232-019-0101-5

48. Usoskin D, Furlan A, Islam S, et al. Unbiased classification of sensory neuron types by large-scale single-cell RNA sequencing. Nat Neurosci. 2015;18(1):145–153. doi:10.1038/nn.3881

49. Schmidt R, Schmelz M, Forster C, Ringkamp M, Torebjiirk E, Handwerker H. Novel Classes of Responsive and Unresponsive C Nociceptors in Human Skin. Vol 15.; 1995.

50. Berkley KJ. Sex differences in pain. Behavioral and Brain Sciences. 1997;20(3):371–380. doi:DOI: 10.1017/S0140525X97221485

